# Exposure to the pesticide tefluthrin causes developmental neurotoxicity in zebrafish

**DOI:** 10.1101/2024.05.28.596249

**Authors:** Fahmi Mesmar, Maram Muhsen, Ibraheem Farooq, Grant Maxey, Jason P. Tourigny, Jason Tennessen, Maria Bondesson

## Abstract

**BACKGROUND:** The insecticide tefluthrin is widely used in agriculture, resulting in widespread pollution. Tefluthrin is a type I pyrethroid characterized by its high persistence in the environment. Understanding the mechanisms of toxicity of tefluthrin will improve its risk assessment.

**OBJECTIVES:** We aimed to decipher the molecular modes of action of tefluthrin.

**METHODS:** Phenotypic developmental toxicity was assessed by exposing zebrafish embryos and larvae to increasing concentrations of tefluthrin. *Tg(mnx:mGFP)* line was used to assess neurotoxicity. Multi-omics approaches including transcriptomics and lipidomics were applied to analyze RNA and lipid contents, respectively. Finally, an *in-silico* ligand–protein docking computational method was used to study a possible interaction between tefluthrin and a protein target.

**RESULTS:** Tefluthrin exposure caused severe morphological malformations in zebrafish larvae, including motor neuron abnormalities. The differentially expressed genes were associated with neurotoxicity and metabolic disruption. Lipidomics analysis revealed a disruption in fatty acid, phospholipid, and lysophospholipid recycling. Protein docking modeling suggested that the LPCAT3 enzyme, which recycles lysophospholipids in the Land’s cycle, directly interacts with tefluthrin.

**CONCLUSIONS:** Tefluthrin exposure causes morphological and neuronal malformations in zebrafish larvae at nanomolar concentrations. Multi-omics results revealed a potential molecular initiating event i.e., inhibition of LPCAT3, and key events i.e., an altered lysophospholipid to phospholipid ratio, leading to the adverse outcomes of neurotoxicity and metabolic disruption.

## Introduction

In the past decades, there has been a considerable rise in neurodegenerative and neurodevelopmental disorders including Alzheimer’s, Parkinson’s, amyotrophic lateral sclerosis (ALS), autism, and attention-deficit/hyperactivity disorder (ADHD). Recent meta-analysis reports demonstrate that these diseases are positively associated with exposure to environmental toxicants such as pesticides (1-5). A class of pesticides suspected to play a role in neurodegenerative and neurodevelopmental disorders is pyrethroid insecticides (6). An association between pyrethroid exposure with ADHD and parent-reported behavioral problems has been reported among US and Canadian children (7, 8). Similar observations were described for rural populations in China. Chen and colleagues demonstrated that prenatal and infant exposures to pyrethroids led to language development delay during childhood (9). Strikingly, Bao *et al*. found that environmental exposure to pyrethroids is associated with an increased risk of overall mortality in the US adult population (10). Although pyrethroids are considered relatively safe due to their low acute mammalian toxicity, Bao and colleagues raise serious concerns about the consequences of chronic exposure on human health (10).

Pyrethroids insecticides are commonly found in consumer products such as pet shampoos, head lice treatments, mosquito sprays, and for lawn and garden insect control. In agriculture, pyrethroids are widely used to treat corn and cotton fields, as well as for treating ectoparasitic infestations in farm animals. These insecticides belong to a large family of synthetic chemicals that can be broadly divided into two groups according to the absence (type I) or presence (type II) of the α-cyano group. Exposure to pyrethroids occurs by direct occupational exposure or by indirect non-occupational exposure through contaminated water, food, or indoor environment. As evidence of their prevalent use, pyrethroid metabolites are detected in people all around the world (11-18). Barr *et al*. showed widespread exposure to pyrethroids in the US (19). They found detectable levels of pyrethroid metabolites in 70% of the analyzed urine samples, with higher exposures in children.

In insects, pyrethroid functions by binding to sodium channels, causing excitatory paralysis followed by death. However in rodents, pyrethroids are known to modulate the activity of several channels of the nervous system, including voltage-gated sodium, chloride, and calcium channels (reviewed in (20)). As fine patterning of neural activity plays a critical role in the maintenance of neuronal survival, aberrant neuronal activity may result in neural apoptosis (21). Several other molecular mechanisms have been proposed to explain pyrethroid-related neurotoxicity including oxidative stress, mitochondrial dysfunction, and neuronal inflammation (22) but further research is required to accurately identify target molecules, especially in nontarget species.

Zebrafish (*Danio rerio*) is increasingly applied for chemical toxicity analysis and drug discovery (23-25). Because the zebrafish nervous system exhibits significant functional similarity and structural homology with other vertebrates including humans (26), zebrafish embryonic and larval stages are popular models for investigating neurotoxicity (27-29). Developmental neuromodulatory effects by pyrethroid exposures have been demonstrated in zebrafish. Spasms appeared after exposures to both type I (e.g., permethrin, resmethrin, and bifenthrin) and type II (e.g., deltamethrin, cypermethrin, and λ-cyhalothrin) pyrethroids in zebrafish larvae (30). Similar to their mammalian counterparts, sodium channels and gamma-aminobutyric acid (GABA)_A_ receptors were the main proposed molecular targets of deltamethrin in zebrafish (31, 32). Additionally, a disruption in the hypothalamus-pituitary-thyroid axis by λ-cyhalothrin, fenvalerate, or permethrin exposure suggests that pyrethroids act as thyroid disruptors (33, 34).

Tefluthrin is a fluorinated highly lipid-soluble type I pyrethroid insecticide. The extensive use of tefluthrin and the environmental persistence of its residue in soil (35) prompted us to investigate whether tefluthrin exposure causes developmental neurotoxicity in zebrafish. Furthermore, we aimed to decipher the molecular mechanisms of tefluthrin-induced neurodevelopmental perturbations by transcriptomic and lipidomic analysis. A potential molecular target for tefluthrin was discovered. A better understanding of tefluthrin toxicity at the molecular level will provide knowledge for an improved risk assessment or for the design of effective countermeasures to oppose exposure-related disease risk.

## Methods

### Chemicals and reagents

PESTANAL® analytical grade tefluthrin (2,3,5,6-tetrafluoro-4-methylbenzyl 3-[(1Z)-2-chloro-3,3,3-trifluoroprop-1-en-1-yl]-2,2-dimethylcyclopropanecarboxylic acid) was purchased from Sigma Aldrich. Tefluthrin stock solutions were prepared in dimethyl sulfoxide (DMSO) (Sigma Aldrich). Freeze-thaw cycles were avoided by storing stock solutions as aliquots at -20 °C until use. The final dilution of working solutions used in all zebrafish experiments were at a DMSO concentration below 0.1%.

### Zebrafish husbandry

Zebrafish were kept in an automated housing system (Aquaneering USA Inc., San Diego, CA). Adult zebrafish (*Danio rerio*) were maintained in tanks supplied with housing water at 28 ± 0.5 °C under 14 h of light and 10 h of dark cycles. Adult fish were fed twice per day with commercial flake food (TetraMin) in the morning and baby brine shrimp (Brine Shrimp Direct) in the afternoon. We used two zebrafish strains: the wildtype NHGRI-1 (Zebrafish International Resource Center, ZIRC) and the transgenic strain *Tg(mnx:mGFP)* (ZIRC). Adult fish were set up for breeding in the late afternoon and the embryos were collected the following morning and maintained in embryo media (E3; 5 mM NaCl, 0.17 mM KCl, 0.33 mM CaCl_2_, 0.33 mM MgSO_4_, pH 7.4) at 28.5°C. All experimental procedures were approved by the Bloomington Institutional Animal Care and Use Committee of Indiana University. We are committed to the highest ethical standards to prevent or minimize pain and distress during the experiment.

### Zebrafish embryo toxicity test

Wild-type zebrafish embryos were used to determine tefluthrin toxicity, largely according to the OECD Fish Embryo Acute Toxicity (FET) Test guidelines (No. 236) (36). Zebrafish embryos were collected in E3 media in a 10 cm plate and placed in the incubator until they reached the shield stage. At 6 hpf, 15 healthy embryos were moved into wells of a 6-well plate containing 5 ml of the exposure media. Zebrafish embryos were exposed to 0.001, 0.01, 0.1, 1, 10, 20, 30, 40, 50, 60, 70, 80, 90, and 100 μM tefluthrin or 0.1% DMSO. Embryos for each concentration were transferred individually in 200 μl exposure media into each well of a 96-well plate. To prevent evaporation, the plates were covered with breathable adhesive seals (ThermoFisher Scientific) and then placed in the incubator at 28.5°C with a light/dark cycle. Static exposure continued without media renewal until developmental toxicity assessments. The embryonic lethality and morphological characteristics were assessed at 72 hpf with a Leica DMi8 microscope equipped with a DMC-4000 color camera. Egg coagulation and heartbeat were used to score for mortality and heart edema, skeletal deformities including spinal curvature, and body position were used as scoring criteria for morphological deformations in accordance with previous studies (37, 38). Each experimental group was tested in three biological replicates to generate concentration-response curves.

### Motor neuron transgenic line and neurotoxicity assessment

Zebrafish transgenic line *Tg(mnx:mGFP)* was used to assess motor neurons and neurite sprouting. At 6 hpf, the embryos were subjected to increasing concentrations of tefluthrin. At 72 hpf, the larvae were imaged using a Leica DMi8 microscope equipped with a DFC9000 camera used to capture the fluorescence of GFP. The images were scored according to the criteria used in Muhsen *et al*. (39).

### Motility analysis in zebrafish embryos

Two dpf NHGRI-1 embryos were exposed for 1 h to 0.1 and 1 μM tefluthrin. Post exposure, they were imaged for 1 minute in a home-built imager using an infrared raspberry pi hq camera with a microscope lens. The movies were analyzed by the software described in the supplementary information, and the total time spent moving (twitching) were summarized/movie.

### RNA extraction, library construction, and sequencing of zebrafish samples

Hatched zebrafish larvae were exposed to two concentrations of tefluthrin (60 nM or 2.3 μM) for 24 h starting at 48 hpf. The exposed larvae were assessed for abnormalities using a stereomicroscope (Leica MZ6). 20–30 larvae with minimal abnormalities were selected and pooled for RNA extraction. Samples were extracted using TRIzol and purified using the Qiagen RNeasy Plus Mini Kit according to the manufacturer’s protocol. The nucleic acid concentration was determined using a microplate reader (Synergy H1; BioTek), and the quality of the RNA was assessed using Agilent 2100 Bioanalyzer system (Agilent Technologies). Samples with high RNA integrity were sequenced by the Center for Genomics and Bioinformatics at Indiana University, Bloomington (https://cgb.indiana.edu). From each sample, 300 ng of total RNA was used to generate polyA-selected libraries following the standard protocol of Illumina strand-specific mRNA library preparation kit protocol. The cleaned adapter-ligated libraries were pooled and loaded on the NextSeq 500 (Illumina) with a 75 high-cycle sequencing module to generate paired-end reads. Sequencing data was submitted to GEO repository (reference number GSE225543).

### Transcriptome assembly, identification of differentially expressed genes, and pathway analysis

The quality of RNA reads was assessed with FastQC (40) and MultiQC (41).Paired-end clean reads were pseudo-aligned and quantified using Salmon v1.6.0 [34] and the D. rerio GRCz11 reference assembly, coding and noncoding transcriptomes retrieved through Ensembl (https://www.ensembl.org), using the chromosomal assembly as the index decoy and the default quantification options. Salmon quantifications were imported into DESeq2 (42) to determine differential gene expression. Cut-off values were selected using log fold change ≥|0.4| and false discovery rate (FDR) < 0.05. Networks, functional analyses, and biological pathways were generated using QIAGEN’s Ingenuity Pathway Analysis (IPA) (https://digitalinsights.qiagen.com/IPA).

### Lipid extraction, lipidomics, analysis of mass spectrometry data and statistic

For lipid profile analysis approximately 25 wildtype zebrafish larvae were exposed to 60 nM tefluthrin from 48 to 72 hpf. Six replicates were analyzed for both control and tefluthrin-treated larvae. Samples were collected on ice in 1.7 ml tubes and then snap-frozen in liquid nitrogen. Lipidomic analysis was performed at the Metabolomics Core Facility at the University of Utah (https://cores.utah.edu/metabolomics/ ). A total of 626 annotated lipids from non-targeted workflow were reported. The methods for lipid extraction and mass spectrometry analysis are described in detail in the Supplementary Information.

For data processing, Agilent MassHunter (MH) Workstation and software packages MH Qualitative and MH Quantitative were used. For lipid annotation, accurate mass and MS/MS matching were used with the Agilent Lipid Annotator library and LipidMatch (43). Results from the positive and negative ionization modes from Lipid Annotator were merged based on the class of lipid identified. Only lipids with relative standard deviations (RSD) less than 30% in quality control samples are used for data analysis. Furthermore, only lipids with background AUC counts in process blanks that are less than 30% of quality control samples are used for data analysis. The raw data were normalized based on the ratio to class-specific internal standards, then to tissue mass and sum before statistical analysis. Multivariate analysis was performed using MetaboAnalyst (44). Statistical models were created for the normalized data after normalizing to sum, logarithmic transformation (base 10), and Pareto scaling. a p-value of < 0.05 and an FC ≥ 1.5 were used for heatmaps or log fold change analysis. For lipid ontology and enrichment analysis LION web was used to assess the changes in the sets of lipids that share a certain property (45).

### Phospholipase A activity

Three dpf NHGRI-1 embryos treated with a range of tefluthrin concentrations were used to investigate phospholipase A activity by live staining with Red/Green BODIPY (Invitrogen) for 2 h, followed by fluorescence microscopy as described above.

### Computational docking

The computational docking was performed using the AutoDock4 workflow (46). BIOVIA Discovery Studio Visualizer was used to visualize the interaction (BIOVIA, D.S. (2015) Discovery Studio Modeling Environment (47). DoGSiteScorer was used for binding site prediction and druggability (48).

### Statistical analysis

GraphPad Prism software (version 9, GraphPad Software Inc) and R statistical software (RStudio) were used to generate the graphs and statistical analysis. One-Way ANOVA and students t-test were used as parametric statistical procedures. Two-Way ANOVA was used to test data with two categorical variables. Nonparametric Kruskal–Wallis test was used to evaluate data that did not fit normality assumption. False discovery rate (FDR) calculations were done to identify differentially expressed genes in the RNA-seq experiment. A distance matrix was computed to overview the similarities and dissimilarities between controls and tefluthrin-treated samples in transcriptomics datasets. Partial least-squares discriminant analysis (PLS-DA) was used to identify initial trends and clusters in lipidomics datasets.

## Results

### Tefluthrin causes morphological malformations in zebrafish larvae

Initially, we identified the concentrations of tefluthrin causing acute toxicity in zebrafish larvae. Concentration-response curves for survival and morphological malformations were determined after treating zebrafish embryos with increasing concentrations of tefluthrin starting from 6 hpf and analyzed at 72 hpf. Zebrafish embryos were statically exposed to 0.001–100 μM tefluthrin or vehicle control.

Tefluthrin-exposed embryos displayed distinct morphological malformations including yolk deformation and lack of mobility (paralyzed in a sidewise body position) at low concentrations (≤ 100 nM). At higher concentrations (≥1 μM), pericardial edema, scoliosis, and severe lordosis were observed (**Fig. 1A**). The EC_50_ for the paralysis phenotype was calculated to 60 nM, whereas for the lordosis phenotype the calculated EC_50_ was 2.3 μM (**Fig. 1B**). Tefluthrin did not cause 100% lethality even at the highest concentration (100 μM) tested. Thus, only a partial mortality curve was generated for tefluthrin (**Fig. 1C**). The results indicate that tefluthrin acts as a potent teratogenic chemical as it caused apical phenotypic changes at low concentrations but did not cause full mortality at any of the tested concentrations. The teratogenic index (TI) for chemicals with such characteristics is estimated to be very high.

**Figure 1.**
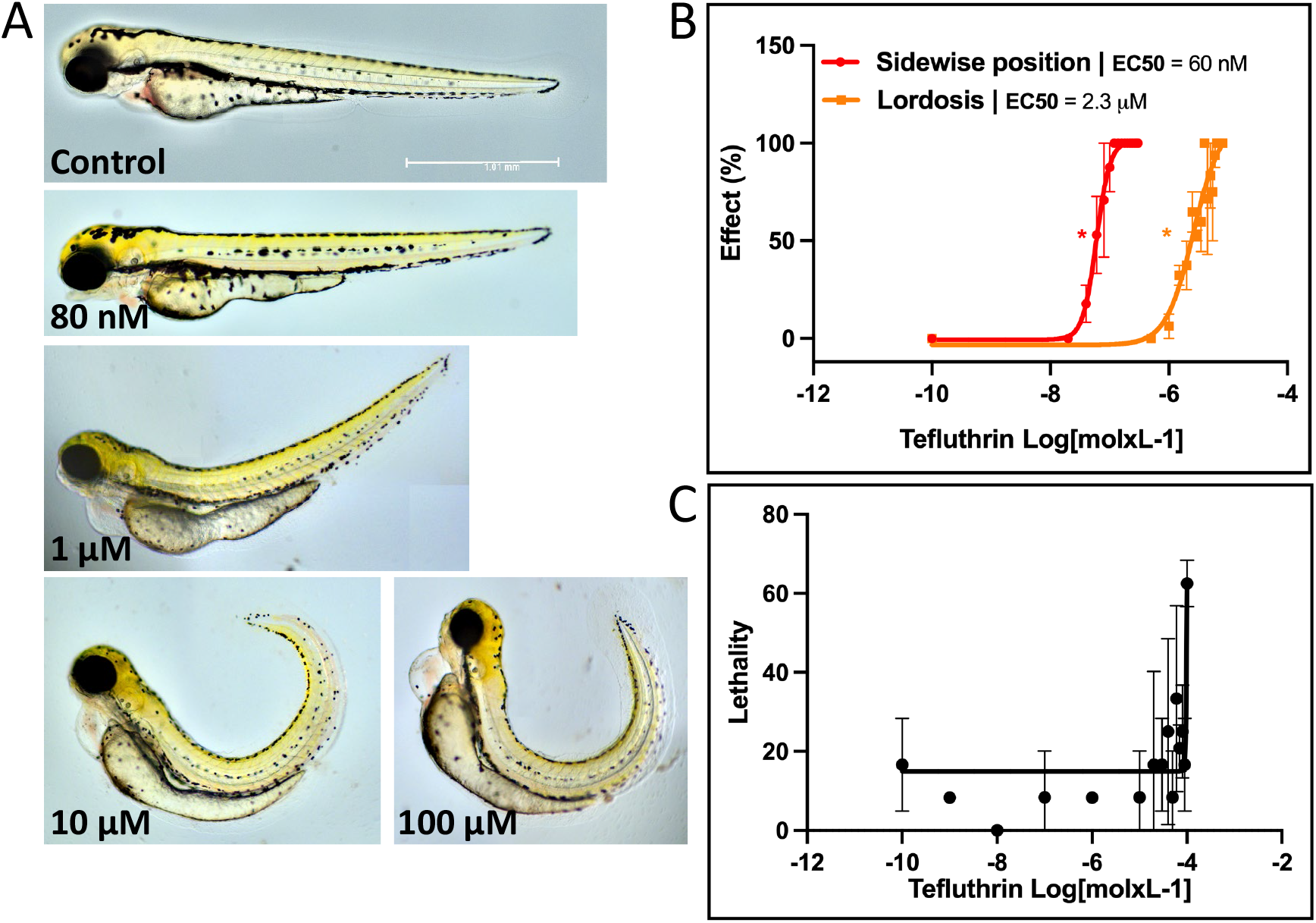
Lethality and teratogenic effects in zebrafish larvae exposed to tefluthrin. **A)** Tefluthrin-treated embryos displayed developmental malformations. **B)** Concentration-response curves of two different apical endpoints, the sidewise position, and lordosis. The EC_50_ of sidewise position was 60 nM and EC_50_ of lordosis was 2.3 μM. **C)** Lethality of tefluthrin exposed larvae. Error bars shown are ± standard deviation of three independent trials. Non-parametric Kruskal–Wallis and Dunn’s multiple comparisons tests were used to calculate p values. *p < 0.05 versus untreated control. Size indicator 1.01 mm.

### Tefluthrin causes secondary motor neuron abnormalities and axonal pathfinding defects in zebrafish larvae

As described above, we observed a paralysis phenotype in larvae laying on their side after tefluthrin exposure. This phenotype may indicate neuromuscular perturbations. To investigate this notion, we utilized the *Tg(mnx:mGFP)* zebrafish transgenic line to examine the effect of tefluthrin on the development of secondary motor neurons and neurite sprouting. *Tg(mnx:mGFP)* embryos were exposed to increasing concentrations of tefluthrin, followed by fluorescence microscopy imaging and axon scoring.

A significant increase in mild to moderate axonal defects was observed in larvae exposed to 100 nM tefluthrin compared to vehicle control (**Fig. 2**). The defects included ectopic or excessive branching, lack of stereotype morphology, axon defasciculation, and innervation in the neighboring myotome, whereas only a few axons showed severe phenotypes. At 250 nM and above, we observed a significant increase in the severe phenotypes including truncated myosepta and complete absence of axons, usually accompanied by excessive branching (**Fig. 3A**). The severity of the axonal perturbations was increased with increasing tefluthrin concentrations (**Fig. 3B**), indicating that exposure interferes with axon development and/or pathfinding leading to neuromuscular impairments in zebrafish.

**Figure 2.**
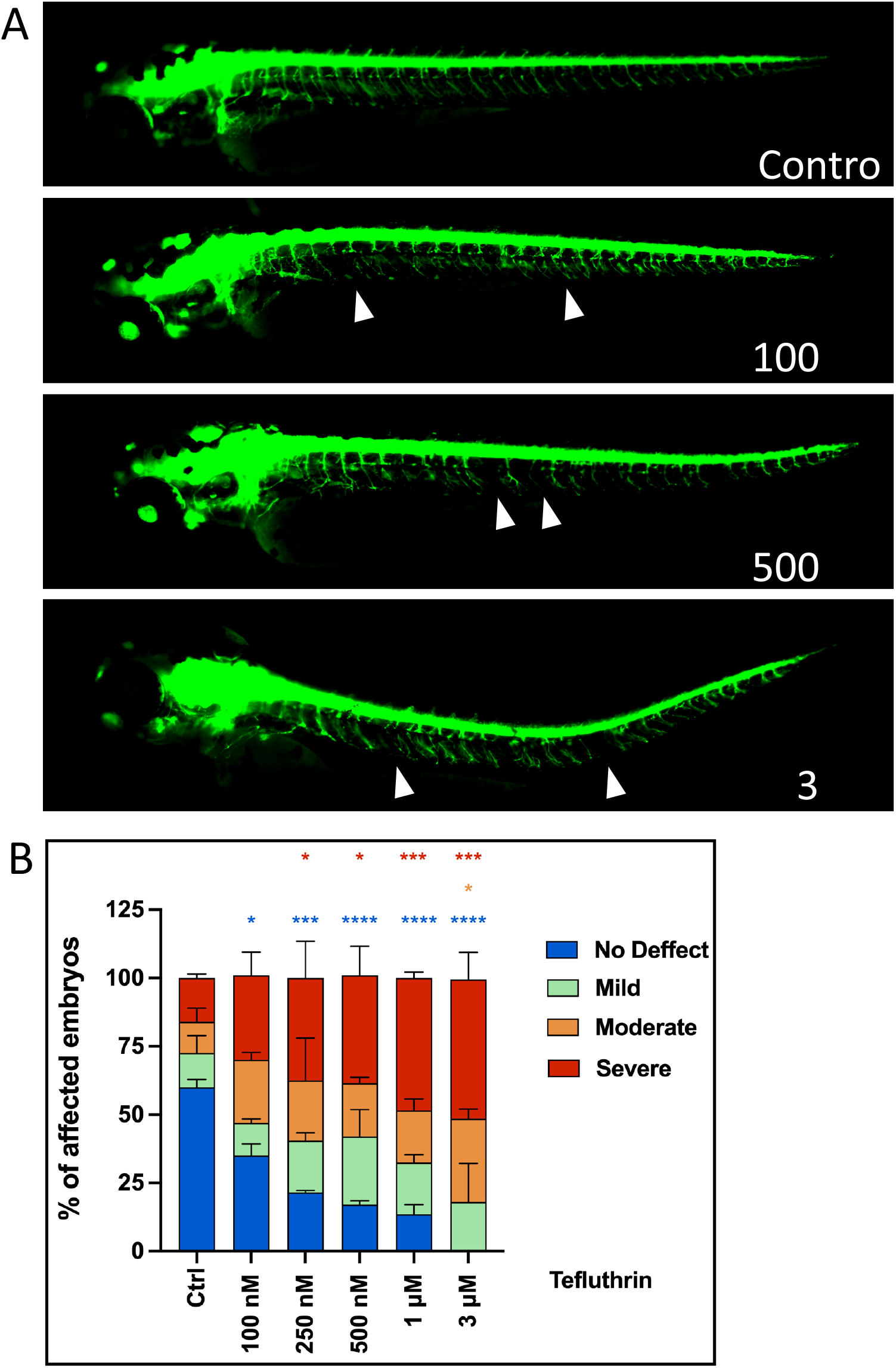
Tefluthrin exposure perturbs secondary motor neuron development in zebrafish larvae. **A)** Lateral view of representative images of 3 dpf *Tg(mnx:mGFP)* larvae. Embryos were treated with increasing concentrations of tefluthrin at 6 hpf (shield stage) and imaged and scored at 72 hpf. Arrowheads point to missing axons or excessive branching. **B)** Motor axon defects were classified as mild, moderate, and severe according to established criteria (39, 49). Two-way ANOVA and Dunnett’s multiple comparison tests were used to calculate p values. Error bars shown are ± standard deviation of three independent trials. *p < 0.05, **p < 0.01, ***p < 0.001, ****p < 0.0001 for sample versus untreated control.

**Figure 3.**
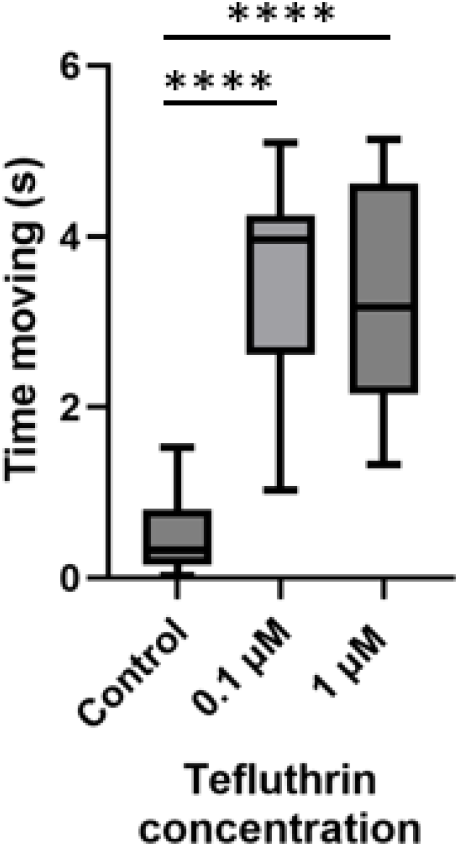
Tefluthrin exposure induces spasm-like behavior. Two dpf zebrafish were exposed to tefluthrin for 1 h, followed by imaging for 1 minute. The time spent moving was analyzed by our generated code (supplementary information) and summarized for each movie. n=17 per concentration and control. **** indicates P<0.0001 by students t-test comparing each treatment to control.

### Tefluthrin acute exposure causes twitching in zebrafish embryos

As shown above, exposure from 6 hpf to 3 dpf results in severe motor neuron perturbations and a paralysis phenotype. However, because other pyrethroids are known to induce hyperactivity, we investigated the consequences of acute exposure on zebrafish embryo motility. Two-days old zebrafish embryos were exposed to 0.1 and 1 μM tefluthrin for 1 h, and motility was imaged subsequently. Both exposure concentrations caused strongly enhanced motility in the form of twitching or a spasm-like behavior (**Fig. 3)**.

### Transcriptomics analysis indicates perturbation of neuronal development, and disruption in carbohydrate and lipid metabolism in tefluthrin-exposed zebrafish larvae

We exposed 48 hpf zebrafish embryos to tefluthrin at 60 nM (corresponding to EC_50_ for paralysis) and 2.3 μM (EC_50_ for lordosis) for 24 h for mRNA sequencing. The low concentration exposure at 60 nM tefluthrin triggered drastic changes in gene expression (**Fig. S1A**) and changed the expression of 6,167 genes (**Fig. S1B**). TP53, HNF4A, CEBPB, CREB1, and FOXO3 (two gene copies in zebrafish, *Foxo3a* and *Foxo3b*) were among the transcription factors that were enriched in response to the low-concentration exposure (**Fig. 4A and Fig. S1C**). Canonical pathway analysis indicated enrichment of cell senescence kinetochore signaling, cell cycle, and DNA repair pathways in treated samples. In addition, an enrichment of the liver X receptor (LXR) and farnesoid X receptor (FXR) as upstream regulators was found; LXR and FXR are nuclear receptors involved in glucose, cholesterol, and lipid metabolism (**Fig. 4B**). Overall, the enrichment of transcription factors that regulate liver metabolic homeostasis, such as LXR, FXR, HNF4A, CEBPB, CREB1, and FOXO3 (50) are indicative of perturbations in the liver transcriptional network. Furthermore, liver toxicity endpoints were among the top enriched toxicity pathways (**Fig. 4D**). Genes that were differentially expressed by tefluthrin were largely associated with carbohydrate and lipid metabolism (**Fig. 4C**). Differentially expressed genes in these pathways (**Fig. S1F**) included *lipca, hsd17b12a*, and *cyp7a1* (**Fig. 4E**). The gluconeogenesis gene *pck1* was upregulated, whereas the glycolysis gene *gck* was downregulated (**Fig. 4F**). In line with the neurodevelopmental malformations observed, IPA confirmed tefluthrin’s neurotoxic effects showing enrichment in the neurological disorder networks (**Fig. S1D and E**).

**Figure 4.**
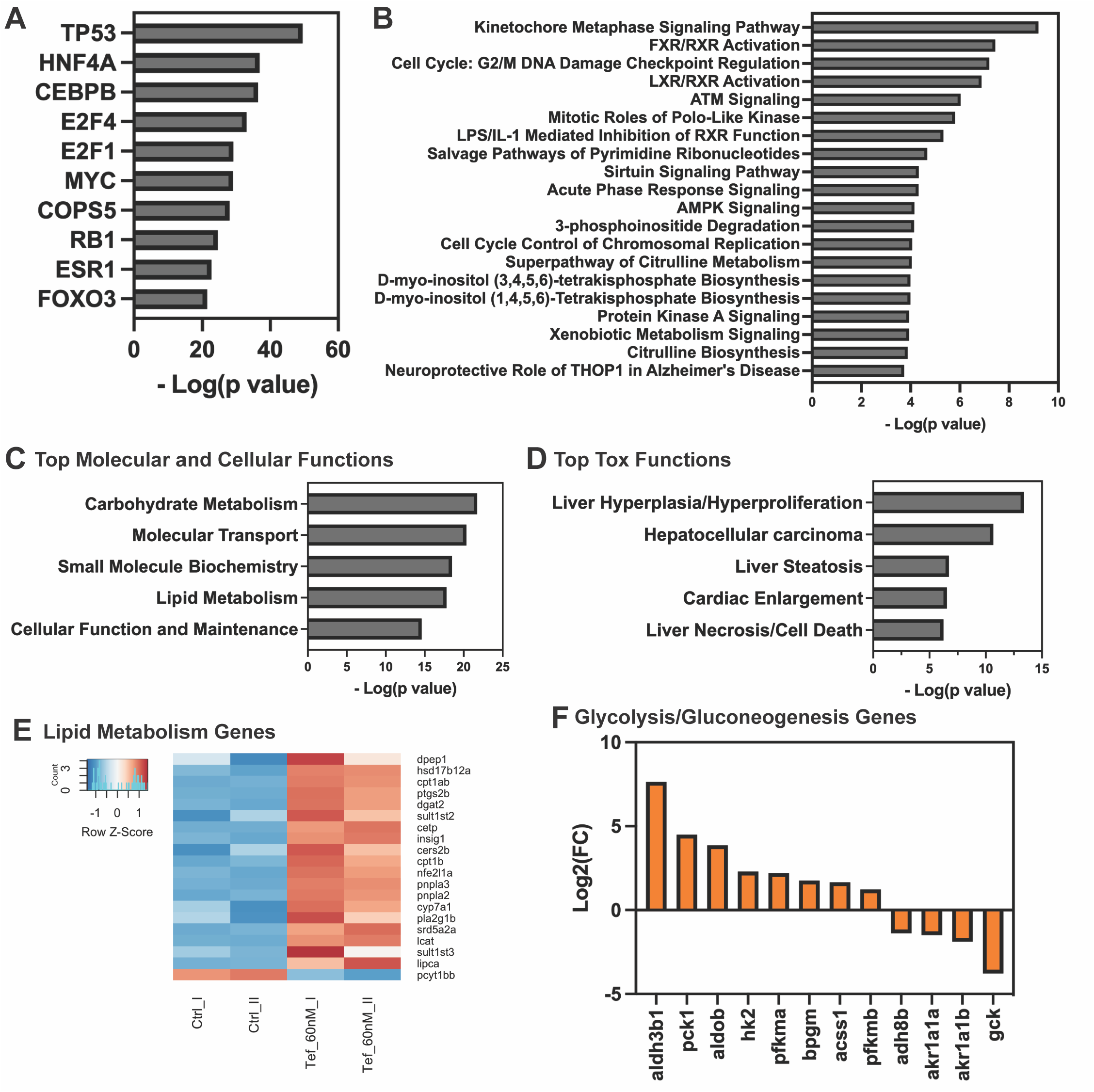
RNA sequencing and pathway analysis of tefluthrin-treated zebrafish larvae. At 48 hpf zebrafish larvae were treated with 60 nM tefluthrin and RNAs were collected at 72 hpf. **A)**. Enriched transcription factors as potential regulators of DEG. **B)** Top 20 canonical pathways by IPA. **C)** Top molecular and cellular functions. **D)** Top toxicity functions. **E)** Genes involved in lipid metabolism pathway. **F)** Genes involved in glycolysis and gluconeogenesis.

For the higher exposure at 2.3 μM, 9,352 genes were differentially expressed (**Fig. S2A**). As for the lower exposure level, exposure to the high concentration of tefluthrin induced enrichment of multiple transcription factors including TP53 (**Fig. S2B**). The activation of TP53-regulated pathways, cell senescence, and autophagy (**Fig. S2C**) indicated that 2.3 μM of tefluthrin was sufficient to induce cell-death-related pathways and toxicity in the zebrafish. Also at this concentration, transcription factor enrichment of HNF4A, CEBPA, and FOXO3-regulated pathways predicted that tefluthrin interferes with metabolic systems as well as causes liver injury. The vulnerability of neurons to tefluthrin was observed as the proliferation of neuronal cells and growth of neurites and nervous tissue were significantly enriched (**Fig. S2D**).

In conclusion, the transcriptomic data suggests that tefluthrin exposure alters gene expression in neuronal cells, interferes with lipid and carbohydrate metabolisms, and causes liver toxicity.

### Lipidomics reveals lipid signatures of tefluthrin exposed zebrafish larvae

To further elucidate the mechanism of tefluthrin toxicity, we applied lipidomics. Zebrafish embryos were exposed to tefluthrin at 60 nM from 48 hpf to 72 hpf. Lipidomics revealed a substantial deviation in the lipid contents in tefluthrin-treated samples (**Fig. S3A**). Lipid ontology analysis showed significant enrichment of several lipid classes including lysoglycerophospholipids and monoacylglycerols with various hydrophilic groups including ethanolamine, choline, and inositol (**Fig. 5A**). To further explore these major lipid categories, we detailed the species with the largest alterations. Various lysophosphatidylethanolamines (LPEs) such as LPE 20:5, 22:5, 18:0, and 16:0, lysophosphatidylcholines (LPCs) such as LPC 18:3, 20:4, 20:2 and many others were among the accumulated lipids in tefluthrin samples, whereas phosphatidylethanolamines (PEs), phosphatidylserines (PSs) and triglycerides (TGs) were reduced (**Fig. 5B and Fig. S3B**). The enrichment in LPE and LPC as well as the decrease in PE, PS, and TG lead us to hypothesize that tefluthrin exposure disrupted phospholipid remodeling in the Land’s cycle, in which two enzymes, lysophosphatidylcholine acyltransferases (LPCAT) and phospholipase A (PLA_2_), control the balance between phospholipids and lysophospholipids.

**Figure 5.**
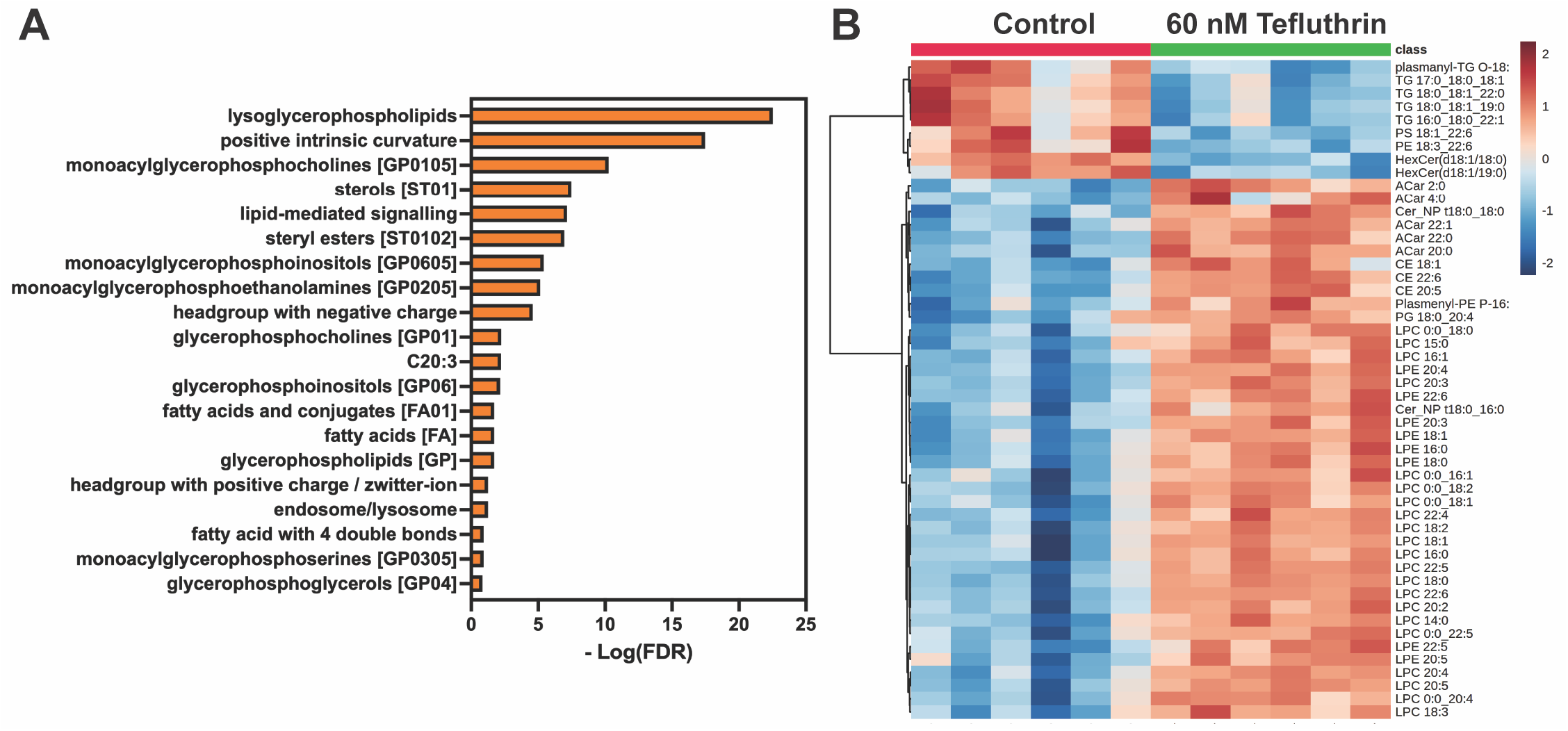
Lipid signatures in zebrafish larvae treated with tefluthrin by non-targeted lipidomics. At 48 hpf zebrafish larvae were treated with 60 nM tefluthrin and then “snap frozen” at 72 hpf (**A)** LION enrichment analysis (lipid ontology) of exposed and control groups. **B)** Heatmap of lipids based on normalized MS intensities in 6 replicates each of control and treatment of the top 50 lipid classes as determined by one-way ANOVA and posthoc analysis.

We were not able to detect any difference in PLA activity after tefluthrin exposure (**Fig. S4**). Thus, the effect of tefluthrin exposure in zebrafish larvae may interfere with LPCATs normal function. After conducting a literature search for LPCAT3 inhibitors we found a single research paper that surveyed thousands of small molecules using a high-throughput screening approach. The authors characterized two chemical hits (HTS-3 and HTS-4) that selectively inhibit LPCAT3 activity (51). Interestingly, there is to some extent structural relatedness between HTS-3, and HTS-4 and tefluthrin (**Fig. S5**).

To examine potential interactions between tefluthrin and LPCAT3 we performed a computational docking study. Utilizing the available crystal structure of *Gallus gallus* LPCAT3 protein (PDB: 7EWT), the *in-silico* docking assay predicated affinity for tefluthrin (**Fig. 6A**). Multiple hydrophobic as well as hydrogen-bonding interactions were formed (**Fig. 6B**) with -9.97 Kcal/mol binding energy and 49.07 nM estimated inhibition constant (ki). Interestingly, tefluthrin was predicted to occupy the binding site with the highest druggability score with a 1072 Å3 pocket size (**Fig. 6C**). Based on this finding, we propose a model of tefluthrin-mediated disruption of lipid homeostasis in the Lands’ cycle, where tefluthrin interferes with phospholipid-fatty acids remodeling through LPCAT3, leading to an increase in lysophospholipid and free fatty acid levels (**Fig. 6D**).

**Figure 6.**
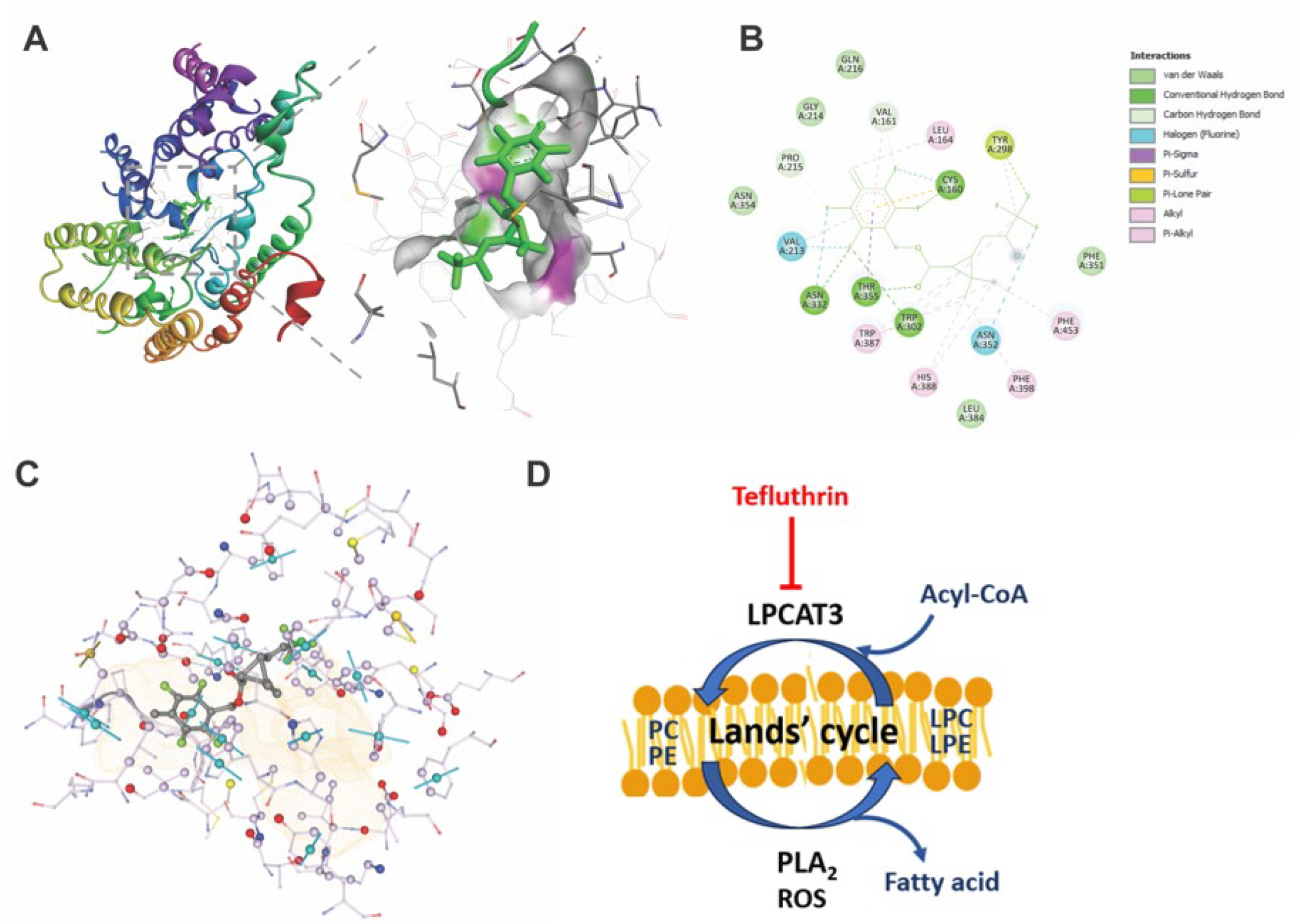
The proposed models for the effect of tefluthrin acting on LPCAT3 to alter lipid homeostasis. **A)** Computational docking of tefluthrin binding to LPCAT3. **B)** Predicated LPCAT3/tefluthrin interactions. **C)** Tefluthrin resides in the largest predicted binding pocket of LPCAT3. **D)** The proposed mechanism for tefluthrin-mediated disruption of lipid homeostasis. Phosphatidylcholine (PC), phosphatidylethanolamine (PE), lysophosphatidylcholine (LPC), lysophosphatidylethanolamine (LPE), lysophosphatidylcholine acyltransferases 3 (LPCAT3), phospholipase A2 (PLA_2_), reactive oxygen species (ROS).

## Discussion

The pyrethroid tefluthrin is widely used as an agriculture insecticide, and due to its fluorinated nature, tefluthrin is persistent with half-life of approximately 1 month. Because of that, more effort is required to assess tefluthrin toxicity and its metabolites, especially in non-target taxa. We here set out to investigate apical phenotypic effects in zebrafish caused by tefluthrin exposure and to decipher its molecular mechanism of action.

Our study demonstrates that exposure to tefluthrin resulted in developmental toxicity in zebrafish larvae. Interestingly, tefluthrin caused severe malformations at substantially lower concentration than lethality, indicating that tefluthrin is a potent teratogen. Our results are in line with those of a large screening study on zebrafish, in which several morphological malformations were described after exposure to tefluthrin (52). Most of these malformations, including yolk sac edema, bent body axis, pericardial edema, malformed jaw, etc., were observed at 6.4 μM exposure. As in our study, tefluthrin exposure did not cause significant mortality (tested up to 64 μM), indicating a high teratogenic index (52). The low level of embryonic mortality in our study could be explained by the presence of the chorion which may protect from exposure to distinct chemicals, thus reducing the uptake and assay sensitivity (53, 54). However, we do not believe that the chorion protected the embryos in our study because enough chemical accumulated in the embryos to produce morphological malformations at low concentrations.

Tefluthrin exhibits neurotoxic effects in both insects and mammals (reviewed in (35)). Additionally, tefluthrin is very toxic to aquatic vertebrates. We demonstrated that whereas a short 1 h exposure caused increased twitching or spasm-like behavior in early embryos, a longer exposure from 6 to 72 hpf caused paralysis and severe lordosis. DeMicco *et al*. observed body axis curvature and spasms in zebrafish larvae treated with six different pyrethroids (not tefluthrin) at relatively low doses (30). Interestingly, when co-treated with the sodium channel antagonist MS222, both the spasm and curvature phenotypes were ameliorated (30), indicating that increased sodium channel signaling results in curvature. To further strengthen the evidence for tefluthrin-induced neurotoxicity, we used *Tg(mnx:mGFP*) zebrafish transgenic line; Mnx being a transcription factor exclusively expressed in the hindbrain and postmitotic spinal cord motor neurons. This model is used to study neuromuscular junction formation and axonal pathfinding (55). Our data indicated that tefluthrin-induced axon toxicity is not selective to the insect nervous system but also causes neurotoxicity in vertebrates. This is in line with a previous study on deltamethrin and permethrin, which also cause developmental neurotoxicity (56). We observed axonal perturbations at very low (100 nM) exposure concentrations in zebrafish. EPA’s acute population-adjusted dose (aPAD) for tefluthrin is 0.005 mg/kg/day (57). With the assumption that 1 kg equals 1 liter, the estimated aPAD is around 12 nM/day, thus, 10 times below the single exposure concentration causing neurodevelopmental perturbations in fish larvae.

We provided mechanistic insights into tefluthrin toxicity through RNA sequencing. Although we in this paper focused our phenotypic assays on neuronal endpoints, bioinformatics identified that the main pathways affected were liver related. Several differentially expressed genes in tefluthrin-treated samples were predicted to be regulated by HNF4A (hepatocyte nuclear factor 4 alpha). *Hnf4a* is highly expressed in the liver and is considered an indicator of hepatic metabolic syndrome (58). HNF4A is important for the normal development and function of the liver hepatocytes and it controls the expression of genes that largely involved in drug metabolism, hepatic lipid metabolism, and gluconeogenesis (59). Additionally, Hnf4a plays a key role in protecting hepatocytes against lipotoxicity (58). Not surprisingly, liver toxicity was the top affected toxicity function. It has been shown recently that *Xenopus laevis* exposure to the pyrethroid cis-bifenthrin results in liver injury associated with impairment in lipid metabolism and NAFLD (60). In the rat model, pyrethroid exposure increases liver damage and tumorigenesis (61, 62).

In addition to HNF4A, RNA-seq analysis identified critical transcription factors that are involved in the transcriptional regulation of metabolism including LXR, FXR CEBPB and FOXO(s). LXR plays a key role in regulating hepatic lipid and carbohydrate metabolisms, as well as lipogenesis and lipolysis in fat tissues (63). FOXO transcription factors are involved in glucose and hepatic lipid metabolism, energy homeostasis, and detoxifying reactive oxygen species (ROS) (64, 65). These factors play a role in liver function and disease (66). Furthermore, FOXO(s) play a critical role in protecting the liver against the development of NAFLD (67) and liver oxidative injury (68). In conclusion, the data indicates that tefluthrin exposure results in liver toxicity resulting in a transcriptomic response to mitigate the toxicity induced.

To further understand the mechanisms and modes of toxic action of tefluthrin, a lipidomics study was carried out on zebrafish. The early larval stage of zebrafish is considered a suitable model to study chemical lipidome interaction because they share conserved lipid metabolic pathways with other vertebrates (69). Moreover, the lecithotrophic criterion of this stage makes them independent from exogenous feeding which can cause considerable experimental variation (70). Lipidomics revealed an increase in lysophospholipids (LPLs) including LPE and LPC. LPLs are lipid molecules that are important signaling molecules for physiological processes, but an imbalance in LPLs levels can also lead to pathological conditions. The lipidomics data revealed that the increase in LPL levels was accompanied by a decrease in phospholipids (PLs). On one hand, PC and PE are targets of peroxidation, which may lead to destabilization of cell and organelle membranes, but on the other hand, LPLs may also destabilize membranes due to their detergent-like character (71). Thus, the levels of PLs and LPLs need to be tightly controlled, which is mediated by LPCAT3 and PLA_2_ in the Land’s cycle (reviewed in (72)).

Our computational protein docking suggested that tefluthrin binds to LPCAT3 activity, and more specifically to residues in the largest predicted binding pocket of LPCAT3. This enzyme is a member of the acyltransferase family that regulates the contents of PL in the liver, adipose tissue (72, 73), and nervous system (74), thus playing an important role in regulating lipogenesis and neuronal lipid homeostasis. Changes in LPCAT activity have been implicated in impaired adipogenesis, non-alcoholic fatty liver disease (NAFLD), and many other pathological conditions (75). For instance, mice lacking *Lpact3* in the liver have hepatocyte lipid accumulation (73). In human liver-derived cell lines, LPE supplementation induces lipid droplet formation (76). In the nervous system, a disruption in LPCAT3 pathway results in neuropathy (74). The addition of LPE supplementation to mouse cortical neuronal cultures stimulates neurite growth (77, 78) , a phenomenon we observed in tefluthrin treated zebrafish. An increase in LPC levels in vivo is associated with neurodegenerative diseases (reviewed in (79)). Furthermore, high LPC mediates motor function defects and neurotoxicity (72). We could not find enough literature supporting possible pyrethroid–LPCAT interaction, but interestingly, Stelzer *et al*. observed an interaction between another pyrethroid, permethrin, with PC by using an *in vitro* liposome system (80, 81). Thus, we propose that tefluthrin directly binds to the LPCAT3 enzyme, or indirectly through its interaction with the LPCAT3 substrate, affecting the phospholipid recycling process. Further studies are needed to test these hypotheses. Finally, our results demonstrated a potential link between metabolic disruption and neurotoxicity. A similar conclusion has been made by Nasri *et al* when they found anilinopyrimidines act as metabolic disruptors and neurotoxicants in zebrafish (82).

In conclusion, the results of the current study provide insight into the mechanisms underlying tefluthrin-related neurotoxicity and induced metabolic dysregulation. In this regard, we propose that disruption in phospholipid recycling is a molecular initiating event (MIE) of tefluthrin exposure, through a series of key events (KE) including motor neuron impairment, and alterations in key metabolic transcription factors leads to the adverse outcomes (AO) of neurotoxicity and metabolic disruption.

## Acknowledgments

We thank the Rural Obesity Project Development Team at Indiana University for fruitful discussions of the study. Metabolomic analysis was performed by John Allen Mack at the Metabolomics Core Facility at the University of Utah. Mass spectrometry equipment was obtained through NCRR Shared Instrumentation Grant 1S10OD016232-01, 1S10OD018210-01A1 and 1S10OD021505-01. RNA sequencing was performed at the Center for Genomics and Bioinformatics at Indiana University Bloomington.

## Supplementary Information

Supplementary Materials and Methods

Figure S1. Pathway analysis of tefluthrin low-concentration exposed zebrafish larvae

Figure S2. Pathway analysis of tefluthrin high-concentration exposed zebrafish larvae

Figure S3. Non-targeted lipidomics of zebrafish larvae

Figure S4. Tefluthrin exposure does not alter PLA_2_ activity

Figure S5. Chemical structure of HTS-3, HTS-4 and tefluthrin

Supplementary Code to analyze embryo movement

## References

1. Bemanalizadeh, M., Khoshhali, M., Goli, P., Abdollahpour, I. and Kelishadi, R., Parental Occupational Exposure and Neurodevelopmental Disorders in Offspring: a Systematic Review and Meta-analysis. Curr Environ Health Rep, 2022. 9(3): p. 406–422.

2. Nabi, M. and Tabassum, N., Role of Environmental Toxicants on Neurodegenerative Disorders. Front Toxicol, 2022. 4: p. 837579.

3. Talbott, E.O., Malek, A.M., Arena, V.C., Wu, F., Steffes, K., Sharma, R.K., Buchanich, J., Rager, J.R., Bear, T., Hoffman, C.A., Lacomis, D., Donnelly, C., Mauna, J., et al., Case-control study of environmental toxins and risk of amyotrophic lateral sclerosis involving the national ALS registry. Amyotroph Lateral Scler Frontotemporal Degener, 2024: p. 1–10.

4. Xu, Y., Yang, X., Chen, D., Xu, Y., Lan, L., Zhao, S., Liu, Q., Snijders, A.M., and Xia, Y., Maternal exposure to pesticides and autism or attention-deficit/hyperactivity disorders in offspring: A meta-analysis. Chemosphere, 2023. 313: p. 137459.

5. Yan, D., Zhang, Y., Liu, L. and Yan, H., Pesticide exposure and risk of Alzheimer’s disease: a systematic review and meta-analysis. Sci Rep, 2016. 6: p. 32222.

6. Arsuffi-Marcon, R., Souza, L.G., Santos-Miranda, A. and Joviano-Santos, J.V., Neurotoxicity of Pyrethroids in neurodegenerative diseases: From animals’ models to humans’ studies. Chem Biol Interact, 2024. 391: p. 110911.

7. Oulhote, Y. and Bouchard, M.F., Urinary metabolites of organophosphate and pyrethroid pesticides and behavioral problems in Canadian children. Environ Health Perspect, 2013. 121(11-12): p. 1378–84.

8. Wagner-Schuman, M., Richardson, J.R., Auinger, P., Braun, J.M., Lanphear, B.P., Epstein, J.N., Yolton, K., and Froehlich, T.E., Association of pyrethroid pesticide exposure with attention-deficit/hyperactivity disorder in a nationally representative sample of U.S. children. Environ Health, 2015. 14: p. 44.

9. Chen, S., Xiao, X., Qi, Z., Chen, L., Chen, Y., Xu, L., Zhang, L., Song, X., and Li, Y., Effects of prenatal and infant daily exposure to pyrethroid pesticides on the language development of 2-year-old toddlers: A prospective cohort study in rural Yunnan, China. Neurotoxicology, 2022. 92: p. 180–190.

10. Bao, W., Liu, B., Simonsen, D.W. and Lehmler, H.J., Association Between Exposure to Pyrethroid Insecticides and Risk of All-Cause and Cause-Specific Mortality in the General US Adult Population. JAMA Intern Med, 2020. 180(3): p. 367–374.

11. Couture, C., Fortin, M.C., Carrier, G., Dumas, P., Tremblay, C. and Bouchard, M., Assessment of exposure to pyrethroids and pyrethrins in a rural population of the Montérégie area, Quebec, Canada. J Occup Environ Hyg, 2009. 6(6): p. 341–52.

12. Dewailly, E., Forde, M., Robertson, L., Kaddar, N., Laouan Sidi, E.A., Côté, S., Gaudreau, E., Drescher, O., and Ayotte, P., Evaluation of pyrethroid exposures in pregnant women from 10 Caribbean countries. Environ Int, 2014. 63: p. 201–6.

13. Hardt, J. and Angerer, J., Biological monitoring of workers after the application of insecticidal pyrethroids. Int Arch Occup Environ Health, 2003. 76(7): p. 492–8.

14. Lu, D., Wang, D., Feng, C., Jin, Y., Zhou, Z., Wu, C., Lin, Y., and Wang, G., Urinary concentrations of metabolites of pyrethroid insecticides in textile workers, Eastern China. Environ Int, 2013. 60: p. 137–44.

15. Panuwet, P., Prapamontol, T., Chantara, S. and Barr, D.B., Urinary pesticide metabolites in school students from northern Thailand. Int J Hyg Environ Health, 2009. 212(3): p. 288–97.

16. Rodzaj, W., Wileńska, M., Klimowska, A., Dziewirska, E., Jurewicz, J., Walczak-Jędrzejowska, R., Słowikowska-Hilczer, J., Hanke, W., and Wielgomas, B., Concentrations of urinary biomarkers and predictors of exposure to pyrethroid insecticides in young, Polish, urban-dwelling men. Sci Total Environ, 2021. 773: p. 145666.

17. Wielgomas, B. and Piskunowicz, M., Biomonitoring of pyrethroid exposure among rural and urban populations in northern Poland. Chemosphere, 2013. 93(10): p. 2547–53.

18. Wu, C., Feng, C., Qi, X., Wang, G., Zheng, M., Chang, X. and Zhou, Z., Urinary metabolite levels of pyrethroid insecticides in infants living in an agricultural area of the Province of Jiangsu in China. Chemosphere, 2013. 90(11): p. 2705–13.

19. Barr, D.B., Olsson, A.O., Wong, L.Y., Udunka, S., Baker, S.E., Whitehead, R.D., Magsumbol, M.S., Williams, B.L., and Needham, L.L., Urinary concentrations of metabolites of pyrethroid insecticides in the general U.S. population: National Health and Nutrition Examination Survey 1999-2002. Environ Health Perspect, 2010. 118(6): p. 742–8.

20. Soderlund, D.M., Clark, J.M., Sheets, L.P., Mullin, L.S., Piccirillo, V.J., Sargent, D., Stevens, J.T., and Weiner, M.L., Mechanisms of pyrethroid neurotoxicity: implications for cumulative risk assessment. Toxicology, 2002. 171(1): p. 3–59.

21. Ikonomidou, C., Triggers of apoptosis in the immature brain. Brain Dev, 2009. 31(7): p. 488–92.

22. Mohammadi, H., Ghassemi-Barghi, N., Malakshah, O. and Ashari, S., Pyrethroid exposure and neurotoxicity: a mechanistic approach. Arh Hig Rada Toksikol, 2019. 70(2): p. 74–89.

23. Bauer, B., Mally, A. and Liedtke, D., Zebrafish Embryos and Larvae as Alternative Animal Models for Toxicity Testing. Int J Mol Sci, 2021. 22(24).

24. Cassar, S., Adatto, I., Freeman, J.L., Gamse, J.T., Iturria, I., Lawrence, C., Muriana, A., Peterson, R.T., Van Cruchten, S., and Zon, L.I., Use of Zebrafish in Drug Discovery Toxicology. Chem Res Toxicol, 2020. 33(1): p. 95–118.

25. Planchart, A., Mattingly, C.J., Allen, D., Ceger, P., Casey, W., Hinton, D., Kanungo, J., Kullman, S.W., Tal, T., Bondesson, M., Burgess, S.M., Sullivan, C., Kim, C., et al., Advancing toxicology research using in vivo high throughput toxicology with small fish models. Altex, 2016. 33(4): p. 435–452.

26. Nelson, J.C. and Granato, M., Zebrafish behavior as a gateway to nervous system assembly and plasticity. Development, 2022. 149(9).

27. de Esch, C., Slieker, R., Wolterbeek, A., Woutersen, R. and de Groot, D., Zebrafish as potential model for developmental neurotoxicity testing: a mini review. Neurotoxicol Teratol, 2012. 34(6): p. 545–53.

28. Kokel, D. and Peterson, R.T., Chemobehavioural phenomics and behaviour-based psychiatric drug discovery in the zebrafish. Brief Funct Genomic Proteomic, 2008. 7(6): p. 483–90.

29. Nishimura, Y., Murakami, S., Ashikawa, Y., Sasagawa, S., Umemoto, N., Shimada, Y. and Tanaka, T., Zebrafish as a systems toxicology model for developmental neurotoxicity testing. Congenit Anom (Kyoto), 2015. 55(1): p. 1–16.

30. DeMicco, A., Cooper, K.R., Richardson, J.R. and White, L.A., Developmental neurotoxicity of pyrethroid insecticides in zebrafish embryos. Toxicol Sci, 2010. 113(1): p. 177–86.

31. Chueh, T.C., Hsu, L.S., Kao, C.M., Hsu, T.W., Liao, H.Y., Wang, K.Y. and Chen, S.C., Transcriptome analysis of zebrafish embryos exposed to deltamethrin. Environ Toxicol, 2017. 32(5): p. 1548–1557.

32. Kung, T.S., Richardson, J.R., Cooper, K.R. and White, L.A., Developmental Deltamethrin Exposure Causes Persistent Changes in Dopaminergic Gene Expression, Neurochemistry, and Locomotor Activity in Zebrafish. Toxicol Sci, 2015. 146(2): p. 235–43.

33. Tu, W., Xu, C., Lu, B., Lin, C., Wu, Y. and Liu, W., Acute exposure to synthetic pyrethroids causes bioconcentration and disruption of the hypothalamus-pituitary-thyroid axis in zebrafish embryos. Sci Total Environ, 2016. 542(Pt A): p. 876–85.

34. Zhang, Q., Zhang, Y., Du, J. and Zhao, M., Environmentally relevant levels of λ-cyhalothrin, fenvalerate, and permethrin cause developmental toxicity and disrupt endocrine system in zebrafish (Danio rerio) embryo. Chemosphere, 2017. 185: p. 1173–1180.

35. Wang, X., Li, H., Wang, S., Martínez, M.A., Ares, I., Martínez, M., Martínez-Larrañaga, M.R., Wang, X., Anadón, A., and Maximiliano, J.E., Tefluthrin: metabolism, food residues, toxicity, and mechanisms of action. Crit Rev Toxicol, 2022. 52(8): p. 664–680.

36. OECD, Test No. 236: Fish Embryo Acute Toxicity (FET) Test. 2013.

37. Selderslaghs, I.W., Van Rompay, A.R., De Coen, W. and Witters, H.E., Development of a screening assay to identify teratogenic and embryotoxic chemicals using the zebrafish embryo. Reprod Toxicol, 2009. 28(3): p. 308–20.

38. Weigt, S., Huebler, N., Braunbeck, T., von Landenberg, F. and Broschard, T.H., Zebrafish teratogenicity test with metabolic activation (mDarT): effects of phase I activation of acetaminophen on zebrafish Danio rerio embryos. Toxicology, 2010. 275(1-3): p. 36–49.

39. Muhsen, M., Youngs, J., Riu, A., Gustafsson, J., Kondamadugu, V.S., Garyfalidis, E. and Bondesson, M., Folic acid supplementation rescues valproic acid-induced developmental neurotoxicity and behavioral alterations in zebrafish embryos. Epilepsia, 2021. 62(7): p. 1689–1700.

40. Andrews, S. A quality control tool for high throughput sequence data. 2010; Available from: https://www.bioinformatics.babraham.ac.uk/projects/fastqc/.

41. Ewels, P., Magnusson, M., Lundin, S. and Kaller, M., MultiQC: summarize analysis results for multiple tools and samples in a single report. Bioinformatics, 2016. 32(19): p. 3047–8.

42. Love, M.I., Huber, W. and Anders, S., Moderated estimation of fold change and dispersion for RNA-seq data with DESeq2. Genome Biol, 2014. 15(12): p. 550.

43. Koelmel, J.P., Kroeger, N.M., Ulmer, C.Z., Bowden, J.A., Patterson, R.E., Cochran, J.A., Beecher, C.W.W., Garrett, T.J., and Yost, R.A., LipidMatch: an automated workflow for rule-based lipid identification using untargeted high-resolution tandem mass spectrometry data. BMC Bioinformatics, 2017. 18(1): p. 331.

44. Xia, J., Sinelnikov, I.V., Han, B. and Wishart, D.S., MetaboAnalyst 3.0--making metabolomics more meaningful. Nucleic Acids Res, 2015. 43(W1): p. W251–7.

45. Molenaar, M.R., Jeucken, A., Wassenaar, T.A., van de Lest, C.H.A., Brouwers, J.F. and Helms, J.B., LION/web: a web-based ontology enrichment tool for lipidomic data analysis. Gigascience, 2019. 8(6).

46. Morris, G.M., Huey, R., Lindstrom, W., Sanner, M.F., Belew, R.K., Goodsell, D.S. and Olson, A.J., AutoDock4 and AutoDockTools4: Automated docking with selective receptor flexibility. J Comput Chem, 2009. 30(16): p. 2785–91.

47. Biovia, D.S. BIOVIA Workbook. 2020; Available from: https://www.3ds.com/products-services/biovia/resource-center/citations-and-references/.

48. Volkamer, A., Kuhn, D., Rippmann, F. and Rarey, M., DoGSiteScorer: a web server for automatic binding site prediction, analysis and druggability assessment. Bioinformatics, 2012. 28(15): p. 2074–5.

49. Carrel, T.L., McWhorter, M.L., Workman, E., Zhang, H., Wolstencroft, E.C., Lorson, C., Bassell, G.J., Burghes, A.H., and Beattie, C.E., Survival motor neuron function in motor axons is independent of functions required for small nuclear ribonucleoprotein biogenesis. J Neurosci, 2006. 26(43): p. 11014–22.

50. Ramatchandirin, B., Pearah, A. and He, L., Regulation of Liver Glucose and Lipid Metabolism by Transcriptional Factors and Coactivators. Life (Basel), 2023. 13(2).

51. Reed, A., Ichu, T.A., Milosevich, N., Melillo, B., Schafroth, M.A., Otsuka, Y., Scampavia, L., Spicer, T.P., and Cravatt, B.F., LPCAT3 Inhibitors Remodel the Polyunsaturated Phospholipid Content of Human Cells and Protect from Ferroptosis. ACS Chem Biol, 2022. 17(6): p. 1607–1618.

52. Truong, L., Reif, D.M., St Mary, L., Geier, M.C., Truong, H.D. and Tanguay, R.L., Multidimensional in vivo hazard assessment using zebrafish. Toxicol Sci, 2014. 137(1): p. 212–33.

53. Henn, K. and Braunbeck, T., Dechorionation as a tool to improve the fish embryo toxicity test (FET) with the zebrafish (Danio rerio). Comp Biochem Physiol C Toxicol Pharmacol, 2011. 153(1): p. 91–8.

54. Panzica-Kelly, J.M., Zhang, C.X. and Augustine-Rauch, K.A., Optimization and Performance Assessment of the Chorion-Off [Dechorinated] Zebrafish Developmental Toxicity Assay. Toxicol Sci, 2015. 146(1): p. 127–34.

55. Zelenchuk, T.A. and Brusés, J.L., In vivo labeling of zebrafish motor neurons using an mnx1 enhancer and Gal4/UAS. Genesis, 2011. 49(7): p. 546–54.

56. Andersen, H.R., David, A., Freire, C., Fernández, M.F., D’Cruz, S.C., Reina-Pérez, I., Fini, J.B., and Blaha, L., Pyrethroids and developmental neurotoxicity - A critical review of epidemiological studies and supporting mechanistic evidence. Environ Res, 2022. 214(Pt 2): p. 113935.

57. EPA, Tefluthrin: Updated Human Health Draft Risk Assessment in Support of Registration Review, O.o.C.S.a.P. Prevention, Editor. 2019, United States Environmental Protection Agency.

58. Pan, X. and Zhang, Y., Hepatocyte nuclear factor 4α in the pathogenesis of non-alcoholic fatty liver disease. Chin Med J (Engl), 2022. 135(10): p. 1172–1181.

59. Lu, H., Crosstalk of HNF4α with extracellular and intracellular signaling pathways in the regulation of hepatic metabolism of drugs and lipids. Acta Pharm Sin B, 2016. 6(5): p. 393–408.

60. Li, M., Lv, M., Liu, T., Du, G. and Wang, Q., Lipid Metabolic Disorder Induced by Pyrethroids in Nonalcoholic Fatty Liver Disease of Xenopus laevis. Environ Sci Technol, 2022. 56(12): p.8463-8474.

61. Deguchi, Y., Yamada, T., Hirose, Y., Nagahori, H., Kushida, M., Sumida, K., Sukata, T., Tomigahara, Y., Nishioka, K., Uwagawa, S., Kawamura, S., and Okuno, Y., Mode of action analysis for the synthetic pyrethroid metofluthrin-induced rat liver tumors: evidence for hepatic CYP2B induction and hepatocyte proliferation. Toxicol Sci, 2009. 108(1): p. 69–80.

62. Price, R.J., Walters, D.G., Finch, J.M., Gabriel, K.L., Capen, C.C., Osimitz, T.G. and Lake, B.G., A mode of action for induction of liver tumors by Pyrethrins in the rat. Toxicol Appl Pharmacol, 2007. 218(2): p. 186–95.

63. Korach-André, M. and Gustafsson, J., Liver X receptors as regulators of metabolism. Biomol Concepts, 2015. 6(3): p. 177–90.

64. Sparks, J.D. and Dong, H.H., FoxO1 and hepatic lipid metabolism. Curr Opin Lipidol, 2009. 20(3): p. 217–226.

65. Zhang, W., Patil, S., Chauhan, B., Guo, S., Powell, D.R., Le, J., Klotsas, A., Matika, R., Xiao, X., Franks, R., Heidenreich, K.A., Sajan, M.P., Farese, R.V., et al., FoxO1 regulates multiple metabolic pathways in the liver: effects on gluconeogenic, glycolytic, and lipogenic gene expression. J Biol Chem, 2006. 281(15): p. 10105–17.

66. Tikhanovich, I., Cox, J. and Weinman, S.A., Forkhead box class O transcription factors in liver function and disease. J Gastroenterol Hepatol, 2013. 28 Suppl 1(0 1): p. 125–31.

67. Dong, X.C., FOXO transcription factors in non-alcoholic fatty liver disease. Liver Res, 2017. 1(3): p. 168–173.

68. Jin, H., Zhang, L., He, J., Wu, M., Jia, L. and Guo, J., Role of FOXO3a Transcription Factor in the Regulation of Liver Oxidative Injury. Antioxidants (Basel), 2022. 11(12).

69. Anderson, J.L., Carten, J.D. and Farber, S.A., Zebrafish lipid metabolism: from mediating early patterning to the metabolism of dietary fat and cholesterol. Methods Cell Biol, 2011. 101: p. 111–41.

70. Fraher, D., Sanigorski, A., Mellett, N.A., Meikle, P.J., Sinclair, A.J. and Gibert, Y., Zebrafish Embryonic Lipidomic Analysis Reveals that the Yolk Cell Is Metabolically Active in Processing Lipid. Cell Rep, 2016. 14(6): p. 1317–1329.

71. Hu, J.S., Li, Y.B., Wang, J.W., Sun, L. and Zhang, G.J., Mechanism of lysophosphatidylcholine-induced lysosome destabilization. J Membr Biol, 2007. 215(1): p. 27–35.

72. Law, S.H., Chan, M.L., Marathe, G.K., Parveen, F., Chen, C.H. and Ke, L.Y., An Updated Review of Lysophosphatidylcholine Metabolism in Human Diseases. Int J Mol Sci, 2019. 20(5).

73. Rong, X., Wang, B., Dunham, M.M., Hedde, P.N., Wong, J.S., Gratton, E., Young, S.G., Ford, D.A., and Tontonoz, P., Lpcat3-dependent production of arachidonoyl phospholipids is a key determinant of triglyceride secretion. Elife, 2015. 4.

74. Ichu, T.A., Reed, A., Ogasawara, D., Ulanovskaya, O., Roberts, A., Aguirre, C.A., Bar-Peled, L., Gao, J., Germain, J., Barbas, S., Masuda, K., Conti, B., Tontonoz, P., et al., ABHD12 and LPCAT3 Interplay Regulates a Lyso-phosphatidylserine-C20:4 Phosphatidylserine Lipid Network Implicated in Neurological Disease. Biochemistry, 2020. 59(19): p. 1793–1799.

75. Wang, B. and Tontonoz, P., Phospholipid Remodeling in Physiology and Disease. Annu Rev Physiol, 2019. 81: p. 165–188.

76. Yamamoto, Y., Sakurai, T., Chen, Z., Inoue, N., Chiba, H. and Hui, S.P., Lysophosphatidylethanolamine Affects Lipid Accumulation and Metabolism in a Human Liver-Derived Cell Line. Nutrients, 2022. 14(3).

77. Hisano, K., Kawase, S., Mimura, T., Yoshida, H., Yamada, H., Haniu, H., Tsukahara, T., Kurihara, T., Matsuda, Y., Saito, N., and Uemura, T., Structurally different lysophosphatidylethanolamine species stimulate neurite outgrowth in cultured cortical neurons via distinct G-protein-coupled receptors and signaling cascades. Biochem Biophys Res Commun, 2021. 534: p. 179–185.

78. Hisano, K., Yoshida, H., Kawase, S., Mimura, T., Haniu, H., Tsukahara, T., Kurihara, T., Matsuda, Y., Saito, N., and Uemura, T., Abundant oleoyl-lysophosphatidylethanolamine in brain stimulates neurite outgrowth and protects against glutamate toxicity in cultured cortical neurons. J Biochem, 2021. 170(3): p. 327–336.

79. Liu, P., Zhu, W., Chen, C., Yan, B., Zhu, L., Chen, X. and Peng, C., The mechanisms of lysophosphatidylcholine in the development of diseases. Life Sci, 2020. 247: p. 117443.

80. Stelzer, K.J. and Gordon, M.A., Interactions of pyrethroids with phosphatidylcholine bilayers: comparisons in liposomal systems exhibiting large or small radii of curvature. Chem Biol Interact, 1985. 54(1): p. 105–16.

81. Stelzer, K.J. and Gordon, M.A., Interactions of pyrethroids with phosphatidylcholine liposomal membranes. Biochim Biophys Acta, 1985. 812(2): p. 361–8.

82. Nasri, A., Valverde, A.J., Roche, D.B., Desrumaux, C., Clair, P., Beyrem, H., Chaloin, L., Ghysen, A., and Perrier, V., Neurotoxicity of a Biopesticide Analog on Zebrafish Larvae at Nanomolar Concentrations. Int J Mol Sci, 2016. 17(12).

