## Supplemental Information for "Exposure to the pesticide tefluthrin causes developmental neurotoxicity in zebrafish"

### Supplementary Information

**Materials and Methods**

***Lipid extraction, lipidomics, and mass spectrometry analysis***

Lipids were extracted using a biphasic solvent system of cold methanol, methyl tert-butyl ether (MTBE) (Fisher Scientific), and PBS/water according to Matyash *et al.* with some modifications (1). 225 µl MeOH with internal standards to were added each sample, then the tubes were briefly vortexed and sonicated for 1 min. EquiSPLASH LIPIDOMIX (Avanti Polar Lipids), cholesterol-d7 (Avanti Polar Lipids), coenzyme Q10 (CoQ10-d9) (Cambridge Isotope Laboratories), and palmitic acid-d31 (Cayman Chemical) were used as lipid standards. Samples were then transferred to 1.4 mm ceramic bead-mill tubes (Qiagen). To recover any remaining sample, 188 µl of PBS was used to wash the initial tubes and then transferred to the corresponding bead-mill tube. Samples were homogenized in the bead-mill tubes for 30 sec. Homogenates were transferred to new 1.7 ml polypropylene tubes (VWR) containing 750 µl MTBE, vortexed briefly, and incubated on ice for 1 h with a brief vortex every 15 min. Following incubation, samples were centrifuged at 15,000 *x g* for 10 min at 4 °C. The organic (upper) layer was collected, and the aqueous (lower) layer was re-extracted with 1 ml of 10:3:2.5 (*v/v/v*) MTBE/MeOH/$H_2O$, briefly vortexed, incubated at room temperature for 15 min, and centrifuged at 15,000 *x g* for 10 min at 4 °C. Upper phases were combined and evaporated to dryness under speedvac. Lipid extracts were reconstituted in 600 µl of 4:1:1 (v/v/v) IPA/ACN/$H_2O$ and transferred to LC-MS vials for analysis. LC-MS-grade solvents were used including isopropanol (IPA) (Honeywell Burdick & Jackson), and acetonitrile (ACN) (Honeywell Burdick & Jackson). Concurrently, a process blank sample and quality control samples were prepared by taking equal volumes from each sample after final resuspension.

A Waters ACQUITYBEH C18 column were used for the separation of lipids. Lipid extracts were separated on an Acquity UPLC CSH C18 column (2.1 x 100 mm; 1.7 µm) (Waters) coupled to an Acquity UPLC CSH C18 VanGuard precolumn (5 × 2.1 mm; 1.7 µm) (Waters) maintained at 65°C connected to an Agilent HiP 1290 Sampler, Agilent 1290 Infinity pump, and Agilent 6545 Accurate Mass Q-TOF dual AJS-ESI mass spectrometer (Agilent Technologies).

Samples were analyzed in a randomized order in both positive and negative ionization modes in separate experiments acquiring with the scan range m/z 100 – 1700. The MS was operated under both positive and negative modes with heated capillaries. For positive mode, the source gas temperature was set to 225 °C, with a drying gas flow of 11 L/min, nebulizer pressure of 40 psig, sheath gas temp of 350 °C, and sheath gas flow of 11 L/min. VCap voltage is set at 3500 V, nozzle voltage 500V, fragmentor at 110 V, skimmer at 85 V, and octopole RF peak at 750 V. For negative mode, the source gas temperature was set to 300 °C, with a drying gas flow of 11 L/min, a nebulizer pressure of 30 psig, sheath gas temp of 350 °C and sheath gas flow 11 L/min. VCap voltage was set at 3500 V, nozzle voltage 75 V, fragmentor at 175 V, skimmer at 75 V and octopole RF peak at 750 V. Mobile phase A composed of ACN:$H_2O$ (60:40, *v/v*) in 10 mM ammonium formate (Sigma; 70221) and 0.1% formic acid (Sigma; 00940) whereas mobile phase B consisted of IPA:ACN:$H_2O$ (90:9:1, *v/v/v*) in 10 mM ammonium formate and 0.1% formic acid. For negative mode analysis, the modifiers were changed to 10 mM ammonium acetate (Sigma; 5.33004). Gradient elution for both positive and negative modes started at 15%

mobile phase B then increased to 30% B over 2.4 min, then increased to 48% B from 2.4 – 3.0 min, then increased to 82% B from 3 – 13.2 min, then increased to 99% B from 13.2 – 13.8 min and held until 16.7 min, and then returned to the initial conditions and equilibrated for 5 min. Flow was 0.4 ml/min throughout, with injection volumes of 7 µl for positive and 10 µl negative mode. Tandem mass spectrometry was conducted using iterative exclusion, the same LC gradient at collision energies of 20 V and 27.5 V in positive and negative modes, respectively.

1. Matyash, V., Liebisch, G., Kurzchalia, T.V., Shevchenko, A. and Schwudke, D., *Lipid extraction by methyl-tert butyl ether for high-throughput lipidomics.* J Lipid Res, 2008. **49**(5): p. 1137-46.

### Figures

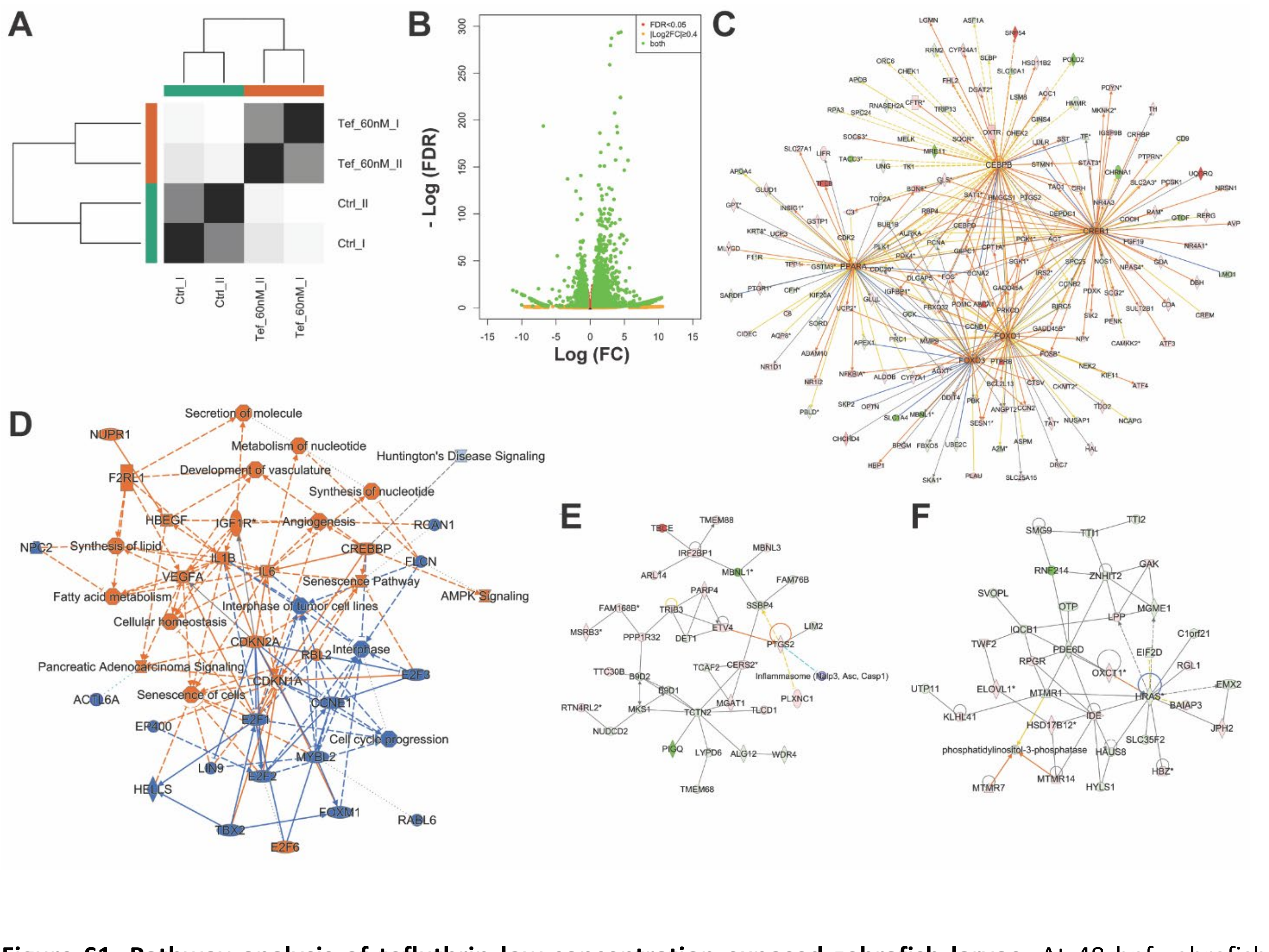


**Figure S1. Pathway analysis of tefluthrin low-concentration exposed zebrafish larvae.** At 48 hpf zebrafish larvae were exposed to 60 nM tefluthrin and RNA was collected at 72 hpf (24 h exposure). **A)** Euclidean distance between samples. **B)** Volcano plot of $\log_2$ (fold change) and $-\log_{10}$ (p adjusted = FDR) of differentially expressed genes (DEG). False discovery rate (FDR) of < 0.05 and |log2 fold change| ≥ 0.4 were used as the cutoff criteria to select DEG. 6167 genes were differentially expressed, 3525 genes were upregulated and 2642 downregulated. **C)** Networks of Cebpb, Creb1 (in zebrafish Creb1a and Creb1b), Ppara (Pparaa and Pparab), Foxo3 and Foxo1 (foxo1a and foxo1b) regulated genes. **D)** Graphical summary of the affected biological, molecular, and cellular functions. **E)** Neurological disorder network. **F)** Lipid and carbohydrate network.

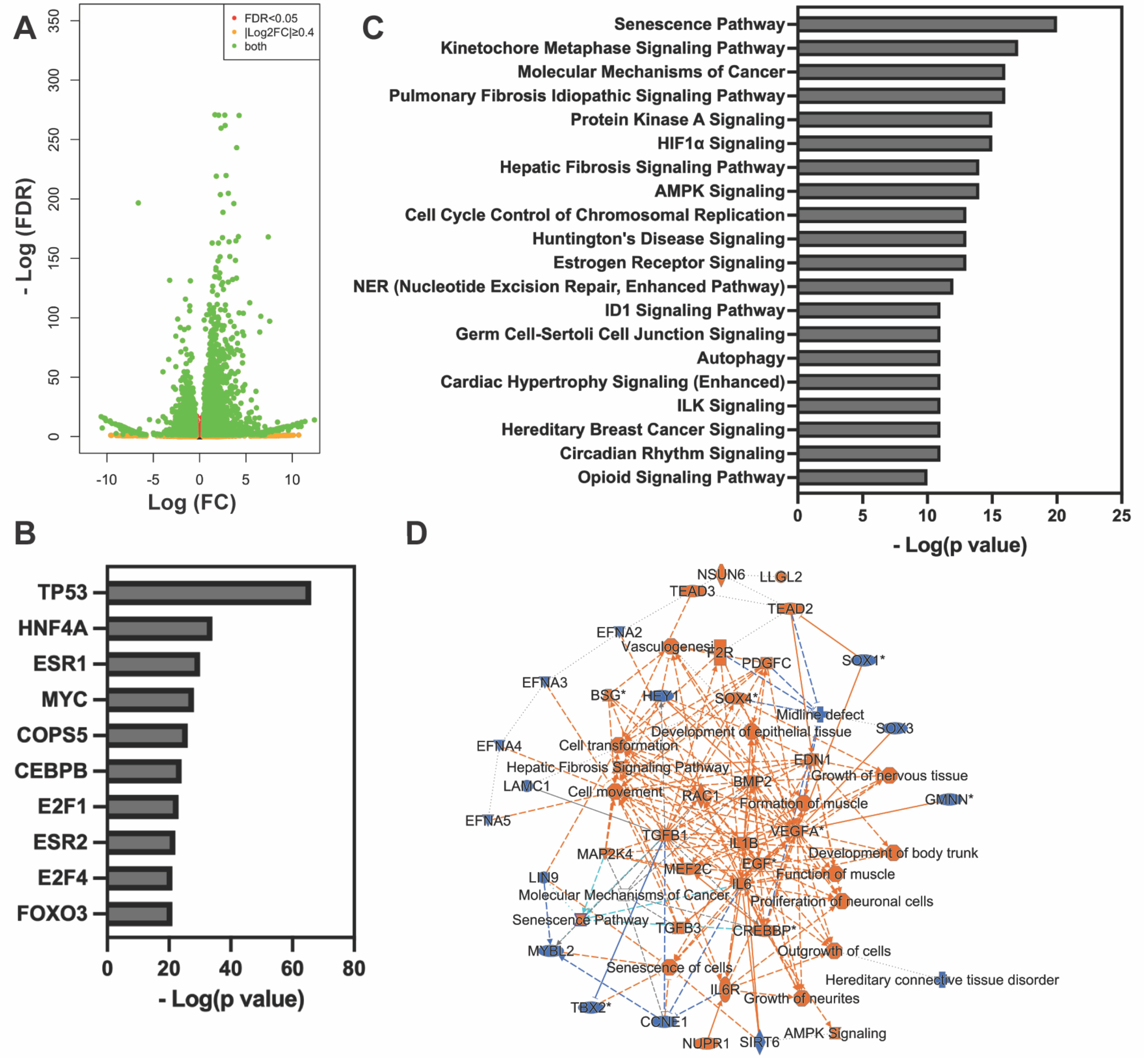


**Figure S2. Pathway analysis of tefluthrin high-concentration exposed zebrafish larvae.** At 48 hpf zebrafish larvae were exposed to 2.3 µM tefluthrin and RNA was collected at 72 hpf (24 h exposure). **A)** Volcano plot of $\log_2$ (fold change) and $-\log_{10}$ (p adjusted = FDR) of differentially expressed genes (DEG). False discovery rate (FDR) of < 0.05 and |log2 fold change| ≥ 0.4 were used as the cutoff criteria to select DEG. 9352 genes were differentially expressed; 5024 genes were upregulated and 4328 downregulated. **B)** Enriched transcription factors predicted to regulate the DEGs. **C)** The top 20 canonical pathways identified by IPA included cell senescence, autophagy, and cell cycle related pathways. **D)** Graphical summary of the affected biological, molecular, and cellular functions.

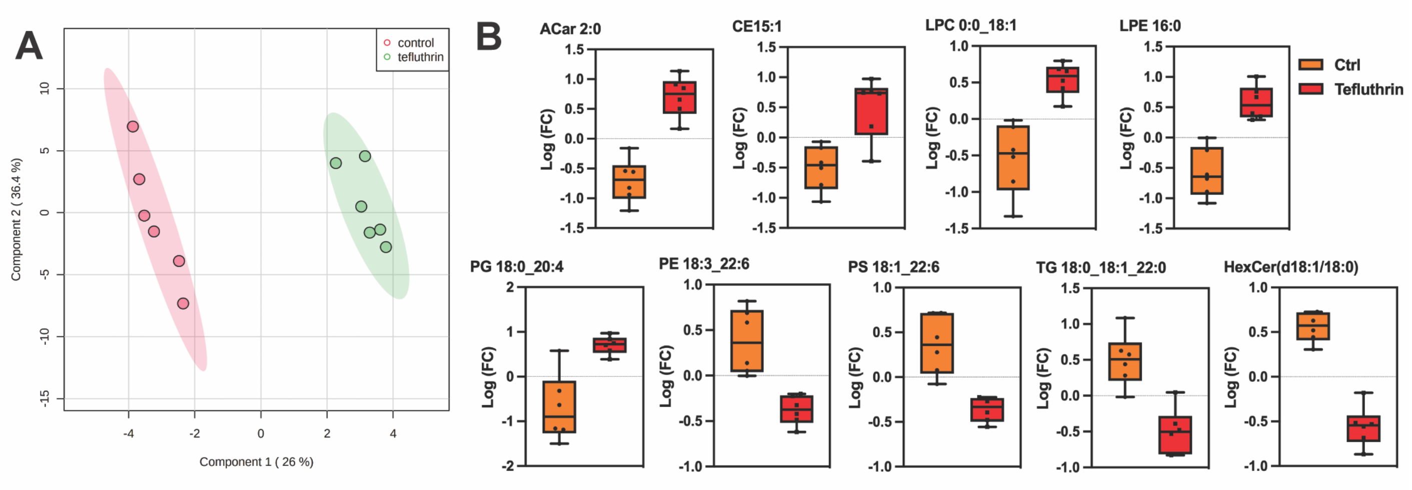


**Figure S3. Non-targeted lipidomics of zebrafish larvae. A)** Partial least-squares discriminant analysis (PLS-DA) of the lipid features in sample treated with vehicle control (DMSO) or tefluthrin of data obtained and merged from positive and negative electrospray ionization modes. **B)** Log (fold change) of normalized MS intensities.

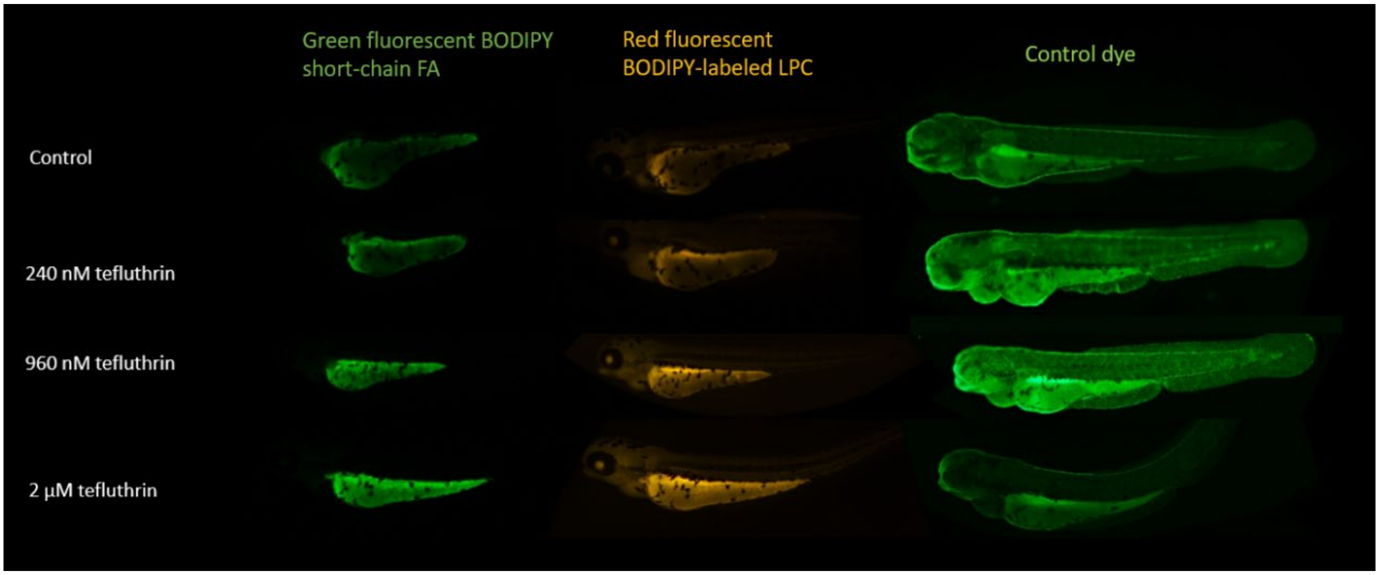


**Figure S4. Tefluthrin exposure** does not alter PLA2 activity. Red/Green BODIPY PC-A2 was added to 3 dpf larvae exposed to tefluthrin for 24 h to score for altered PLA2 activity. No alteration in the green to red ratio was detected, indicating that PLA2 activity was not affected by tefluthrin exposure. The stain mainly accumulated in the yolk.

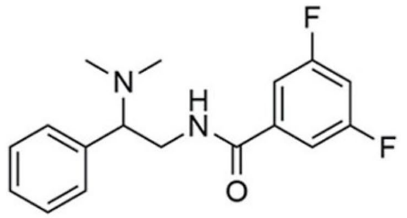

**HTS-3**

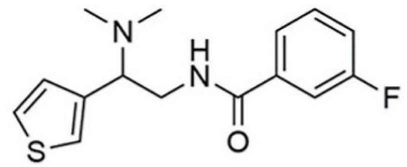

**HTS-4**

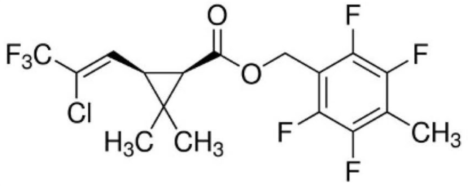

**Tefluthrin**

**Figure S5. Chemical structure of HTS-3, HTS-4 and tefluthrin.** Structures of HTS-3 and HTS-4 are from Reed *et al.* (2). The three compounds share the fluorotoluene moiety.

2. Reed A, Ichu TA, Milosevich N, Melillo B, Schafroth MA, Otsuka Y, Scampavia L, Spicer TP, Cravatt BF. LPCAT3 Inhibitors Remodel the Polyunsaturated Phospholipid Content of Human Cells and Protect from Ferroptosis. ACS Chem Biol. 2022 Jun 17;17(6):1607-1618.
